# Contribution of default mode network to game and delayed-response task performance: power and connectivity analyses of theta oscillation in the monkey

**DOI:** 10.1101/2023.03.19.533320

**Authors:** Tohru Kodama, Takashi Kojima, Yoshiko Honda, Takayuki Hosokawa, Akihiro Karashima, Masataka Watanabe

## Abstract

Neuroimaging studies have demonstrated the presence of a default mode network (DMN) which shows greater activity during rest, and an executive network (EN) which is activated during cognitive tasks. DMN and EN are thought to have competing functions.

However, recent studies reported that the two networks show coactivation during some cognitive tasks. To clarify how DMN works and how DMN interacts with EN for cognitive control, we recorded EEG activities in the medial prefrontal (anterior DMN: aDMN), posterior cingulate/precuneus (posterior DMN: pDMN), and lateral prefrontal (EN) areas in the monkey. As cognitive tasks, we employed a monkey-monkey competitive video game (GAME) and a delayed-response (DR) task. We focused on theta oscillation because of its importance in cognitive control. We also examined theta band connectivity among the three network areas using the Granger causality analysis.

DMN and EN were found to work cooperatively in both tasks. In all the three network areas, we found GAME-task-related, but no DR-task-related, increase in theta power from the resting level, maybe because of the higher cognitive demand associated with the GAME task performance. The information flow conveyed by the theta oscillation was directed more to aDMN than from aDMN for both tasks. The GAME-task-related increase in theta power in aDMN is supposed to be supported by more information flow conveyed by the theta oscillation from EN and pDMN.

## 1. Introduction

Human neuroimaging studies have demonstrated the presence of a default mode network (DMN) and an executive network (EN) [1]. The former consists mainly of the medial prefrontal cortex, posterior cingulate/precuneus areas, and inferior parietal cortex. The latter consists mainly of the lateral prefrontal and posterior parietal cortices. EN is activated while DMN is deactivated during cognitive tasks [1]. DMN and EN are thought to have competing functions. Recent studies have reported that the two networks are not necessarily competing, and instead they actually show coactivation during some cognitive tasks [2]. Neural mechanisms regarding how DMN contributes and how DMN interacts with EN for cognitive control are poorly clarified. To search for these mechanisms, EEG/EMG activities, especially theta (θ) band oscillations that are considered to be important for cognitive control [3], have been investigated. An MEG study [4] indicated that directed attention was associated with an increase in θ oscillation both in DMN and EN. Kam et al. [5] recorded intracranial EEG activities from the DMN and EN areas in presurgical patients, and found that enhanced θ band connectivity between the DMN and EN was a core electrophysiological mechanism that underlies internally directed attention.

We recorded EEG activities from both the DMN and EN regions in the monkey, in relation to its cognitive (video game and delayed-response) task performance as well as the resting state. As DMN, we selected the medial prefrontal (anterior DMN: aDMN) and posterior cingulate/precuneus (posterior DMN: pDMN) areas. As EN, we selected the dorsolateral prefrontal area. We focused on event-related changes in θ band oscillation and examined θ band connectivity among the three network areas by using the Granger causality analysis.

## 2. Material and Methods

### 2.1. Subjects

Two Japanese monkeys (*Macaca fuscata*) (Monkey Y, female weighing 5.5 kg, and Monkey H, male weighing 6.5 kg), were used for this experiment. All experiments were conducted in accordance with the US National Research Council’s Guide for the Care and Use of Laboratory Animals and were approved by the ethics committee of our institute.

### 2.2. Task training

The monkeys were trained on two kinds of cognitive task which were detailed in our previous papers [6,7]. For both tasks, the monkey was seated on a primate chair without head restraint, facing a computer display, joystick, and start button that were vertically arranged (Fig. 1A).

**Fig. 1.**
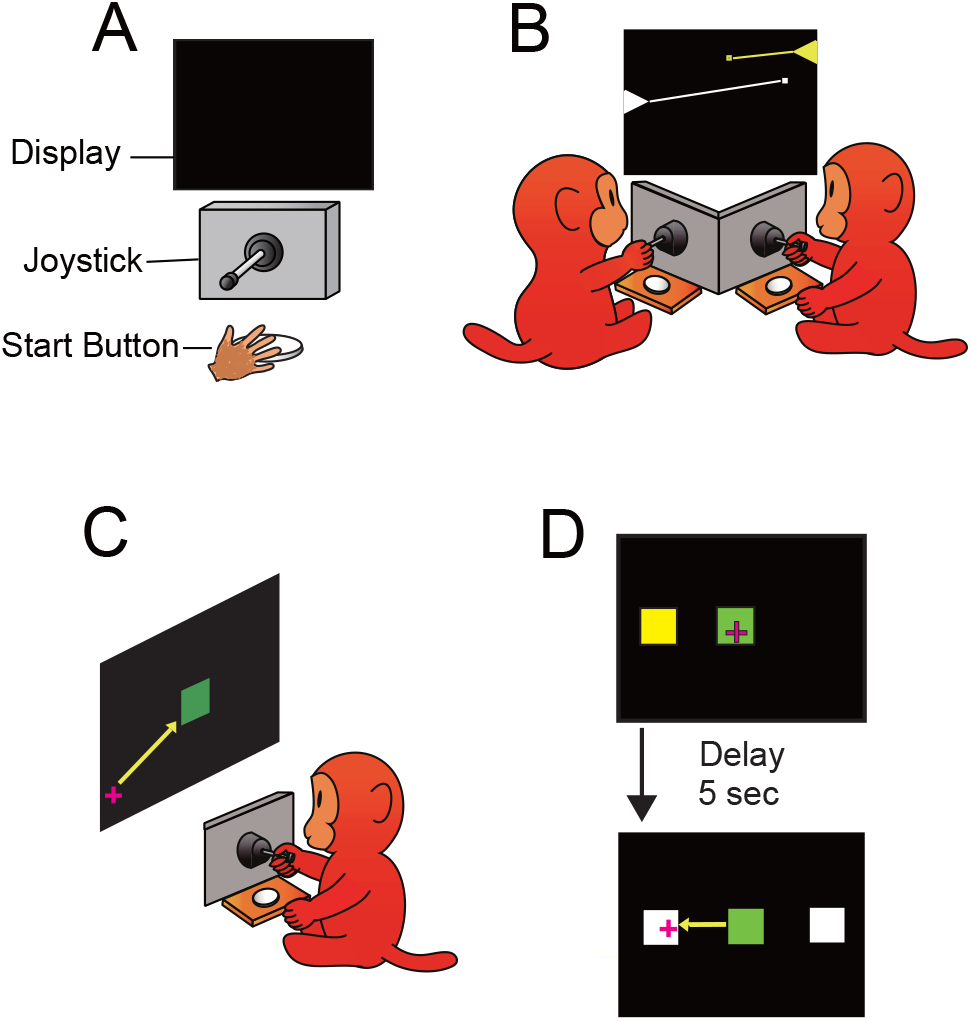
Game and DR Tasks. Task descriptions of the competitive video game and delayed-response task. (A) Experimental setup. (B) The lines on the display represent the trajectories of the bullets and did not appear in the actual game. (C, D) The yellow arrows in C and D did not appear in the actual task. (Color should be used)

In the video game (GAME) task, the two monkeys competed against each other (Fig. 1B). When two colored (white and yellow) triangles appeared on the left and right sides of the display, the monkeys tilted the joystick and shot at the triangle (target) on the other side. The color of the triangle was fixed for each monkey: yellow for monkey Y, and white for monkey H. The monkey that made the first successful hit was the winner and obtained a drop of grape juice (0.3 ml) as a reward.

In the delayed-response (DR) task, which is a typical working memory task, the animal was required to move the plus-mark cursor to the center square (green) by manipulating the joystick (Fig. 1C). Then, a yellow square cue was presented for 0.5 s either on the right or left side of the display. There was then a delay period of 5 s. When two white squares were presented as a go signal on both the right and left sides of the display, the animal could obtain a reward by moving the cursor to the white square where the yellow cue had been presented (Fig. 1D).

### 2.3. EEG recording

After both monkeys were well trained, they were surgically prepared for the EEG recordings under sodium pentobarbital anesthesia (20 mg/kg body weight, i.v.). Concentric bipolar electrodes (silver needles of 0.4 mm diam. insulated with cashew except at recording sites) with the distance between the two recording sites being 0.1mm, were implanted in the DMN and EN areas, based on MRIs (whole-brain coverage, 2 mm slice thickness; Sonata 1.5 T; Siemens) that were obtained before the experiment. The tip of the electrode was aimed to be located in the middle of the layer of the cortical gray matter. For each monkey, 4 electrodes were implanted in EN, 6 electrodes in aDMN, and 4 electrodes in pDMN (See Fig. 4 for the areas of EN, aDMN, and pDMN).

**Fig. 2.**
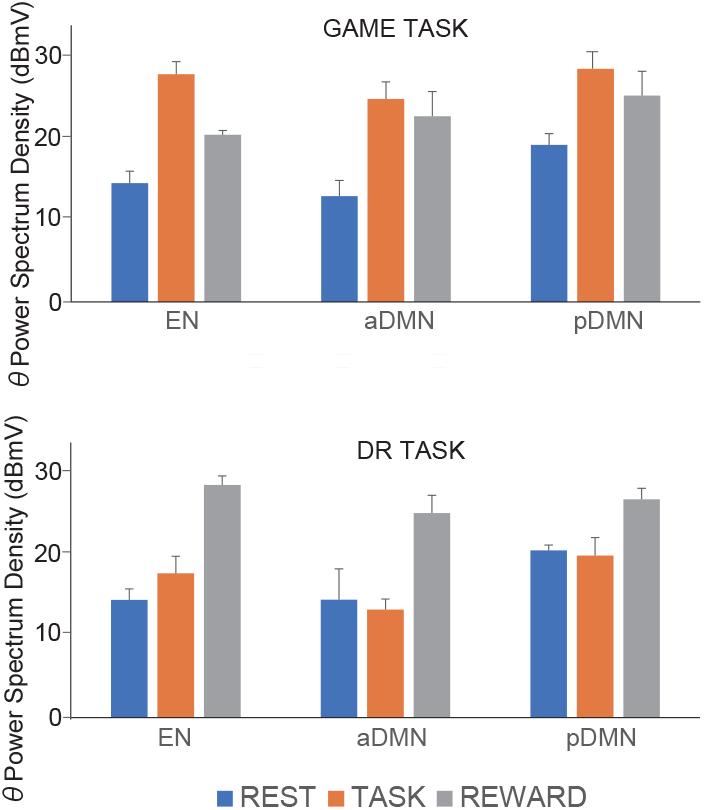
θ power during the REST, TASK, and REWARD periods for the EN, aDMN, and pDMN areas. The θ power is shown separately for the GAME task (upper) and DR task (lower) during the REST (blue), TASK (GAME or DR) (red), and REWARD (grey) periods for the EN, a-DMN, and p-DMN areas. Error bars indicate the SEM. (Color should be used)

**Fig. 3.**
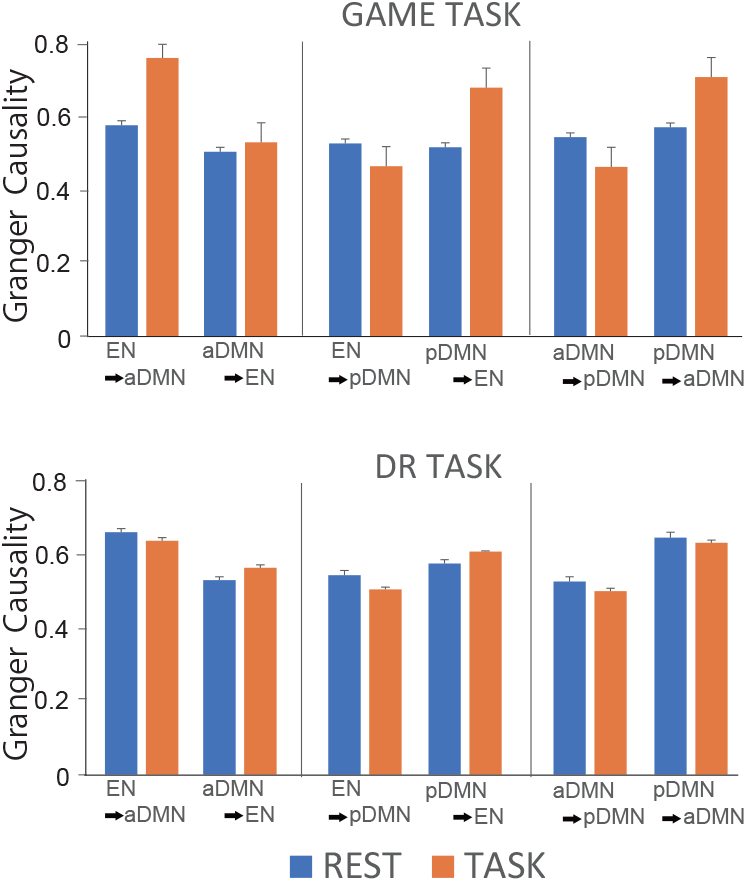
Granger causality (GC) data for GAME and DR tasks. Granger causality (GC) is shown separately for the GAME task (upper) and DR task (lower). GC for the information flow from EN to aDMN, aDMN to EN, EN to pDMN, PDMN to EN, aDMN to pDMN, and pDMN to aDMN is shown separately for the REST period (blue) and the TASK period (red). Error bars indicate the SEM. (Color should be used)

**Fig. 4.**
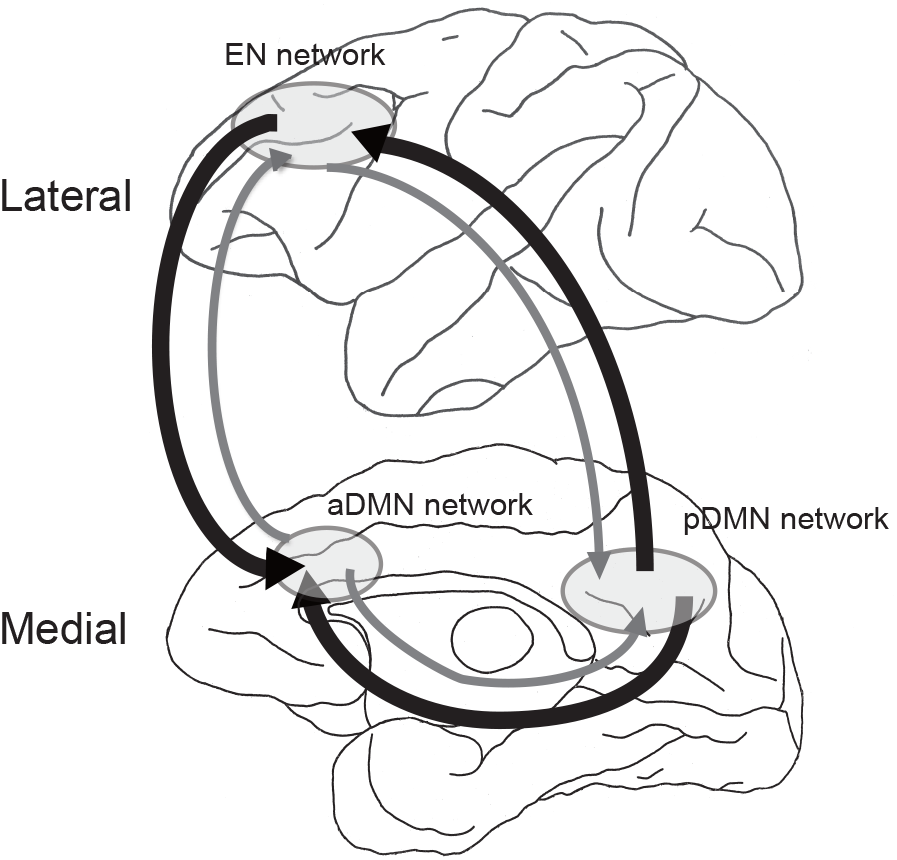
Schema of information flow among the three networks. Thick lines indicate a larger amount of information flow than the thin lines.

In the experiment, the monkey was partially water deprived. In a daily session, the monkey performed either a GAME or DR task. During the resting period, the monkey sat on the monkey chair without task performance in the dark and quiet experimental room. The rest and task periods were alternated with each duration being 30 min. Each monkey performed only one task for an entire day. On the day of GAME, the two monkeys were simultaneously moved, while on the day of DR, each monkey was separately moved, to the experimental room. The monkey obtained fluid only by performing the task during the experimental days, while they were allowed to access free water during the weekend. Monkey pellets were available anytime, and supplemental vegetables and fruits were given after the experiment in the home cage.

### 2.4. EEG data analysis

EEG signals from the 14 bipolar electrodes were recorded at a sampling rate of 1 kHz using a Cerebus data acquisition system (Blackrock, UT, USA). All the data analyses were performed with Matlab code (Mathworks, Natick, MA) using the FieldTrip toolbox (https://www.fieldtriptoolbox.org/). The recording data were first down-sampled to 200 Hz, then, the differential activity between the bipolar electrode was calculated as a local cortical field potential (LFP). Fast Fourier transformation (FFT) was performed following hanning tapering of the EEG data to obtain Power Spectrum Density (PSD) data. The transition of spectra was calculated by shifting the data window in 100ms steps with overlapping. The analysis window for the task period covered 1 s from the time of the “target presentation” in the GAME task, and covered 6 s from the “spatial cue presentation” in the DR task. For both tasks, the analysis window for the reward period was 0.5 s after its delivery. Data only on winning trials in the GAME task and only on correct trials in the DR task were analyzed. The 1 s window in the GAME task and the 6 s window in the DR task were defined as the ‘TASK period’, and the 0.5 s after the reward delivery was defined as the ‘REWARD period’. The data during rest were consecutively segmented into artifact-free epochs of 7000 ms (‘REST period’) without overlapping.

To represent θ oscillation data for each of the three behavioral situations at each electrode point, the mean power of θ oscillation was obtained for each day from 100 representative samples selected usually from the middle time of the second REST and TASK periods. The averaged PSD values of θ spectral band (3-8 Hz) obtained from the 10 experimental days were used as the representative PSD values.

### 2.5. Granger causality

Directionalities of the information flow among EN, aDMN, and pDMN for the periods of REST and TASK were analyzed using Granger causality (GC) in the frequency domain of θ band [8]. GC was calculated using GC toolbox, developed by Seth [9]. Six (two EN-aDMN, two EN-DMN, and two aDMN-pDMN) combinations of GC values in each experimental situation were used for ANOVA analyses.

## 3. Results

### 3.1. Behavior

The monkey performed usually more than 1000 rewarded trials obtaining 300 ml of fruit juice every day on both the GAME and DR days. In the GAME task, both monkeys won the game evenly. In the DR task, both monkeys attained more than 98% correct performance in a daily session. Behavior during REST differed between the two monkeys: while Monkey Y was calm and motionless, Monkey H was restless moving its head and hands on the monkey chair, especially during REST on GAME days when the two monkeys were in the same room. Its θ oscillation data were noisy and enough reliable data were not obtained from this monkey. Thus, we describe here only data obtained from Monkey Y in which reliable data without noise were obtained from 9 points (4 EN, 3 a-DMN, and 2 p-DMN points).

### 3.2. Task related θ power changes in EN, aDMN, and pDMN areas

Figure 2 indicates θ power (in dBmV) during the REST, TASK (GAME or DR), and REWARD periods for the EN, aDMN, and pDMN areas separately for the GAME day and DR day. We applied a three-way ANOVA of task type (GAME and DR) by network (EN, aDMN, and pDMN) by behavioral situation (REST, TASK, and REWARD periods). The statistically significant level was set at P<0.05. There were significant main effects of network [F (2,24) =5.575, P<0.05] and behavioral situation [F (2,24) =78.422, P<0.0001], but not of task type. There was no significant difference in θ power between the EN and the two DMN areas while the θ power was significantly higher in pDMN than in aDMN. There were significant differences in θ power among the three behavioral situations with the power being the highest during the REWARD period and the lowest during the REST period. There was a significant interaction between the task type and behavioral situation [F (2,24) =58.855, P<0.0001]. Post hoc tests using Ryan’s method [10] indicated significant differences in θ power between the two task types during the TASK and REWARD periods, but not during REST. For both task types, there were significant differences in θ power depending on the behavioral situation; in relation to the GAME task, the θ power was the highest during GAME, and the lowest during REST, and in relation to the DR task, the power was larger during the REWARD period than during the REST and TASK periods while the power was not significantly different between the TASK and REST periods. A reward-related increase in θ power from the resting level was observed in both tasks.

### 3.3. Information flow by θ wave among the EN, aDMN, and pDMN

Figure 3 shows the results of Granger causality analyses applied on the amount of information flow by θ wave among the three networks (EN, aDMN, and pDMN) during the REST and TASK periods. A three-way ANOVA (factors of task type, direction of information flow, and behavioral situation (REST vs. TASK)) revealed significant main effects of direction of information flow [F (1, 24) = 52.294, p<0.0001] and of behavioral situation [F (1, 24) = 17.224, p<0.0001], but not of task type. The interactions between the factors of task type and direction of information flow, between the factors of the direction of information flow and behavioral situation, and among the three factors were significant (P<0.0001). As for the direction of information flow, the amount of information flow from EN to aDMN, that from p-DMN to EN, and that from pDMN to aDMN, was significantly larger than the amount of opposite direction of information flows.

Post-hoc analyses using Ryan’s method [10] indicated that for the TASK periods (both GAME and DR), the amount of information flow from EN to aDMN, that from pDMN to EN, and that from pDMN to aDMN was larger than the amount of opposite direction of information flows, although for the REST period, no such clear tendency was observed (Fig. 3). During the GAME task, the amount of information flow for all directions combined, was significantly larger during the TASK than during the REST period. Furthermore, the amount of information flow was significantly larger during GAME than during REST for the direction from EN to aDMN, that from pDMN to EN, and that from pDMN to aDMN, but no such tendency was observed for the opposite directions of information flow. During the DR task, there was no significant difference in the amount of information flow between the TASK and REST periods for any direction among the three networks (Fig. 3).

## 4. Discussion

In our previous neuroimaging (PET) study on monkeys [11], we found a task-related decrease in blood flow in the DMN areas. Since human neuroimaging studies have indicated negative correlations between the θ power and fMRI signal in the DMN [12], we might have found a task-related increase in θ power in DMN and decrease in EN. However, as described before, EN and DMN are not necessarily anticorrelated [2]. We actually did not observe anticorrelation, but found similar activities in θ oscillation between the EN and DMN.

We found a significant increase in θ power from the rest level in relation to the GAME task performance in both the DMN and EN. The θ activity in aDMN is shown to support general attentional or executive function [13], and is reported to be greater when the task becomes more demanding [14]. To win the competitive game, the monkey was required, to observe which locus the target appeared in, to manipulate the joystick according to which direction the bullet should be shot, and to shoot the bullet as fast and as accurately as possible. During the DR task, the monkey was concerned mainly with retaining spatial information in working memory without paying attention to external stimuli except at the time of the cue presentation. For the well-trained monkey, this operation may not be so cognitively demanding, and the monkey performed the DR task almost perfectly. Thus, the increase in θ power during the GAME task from the rest level may be caused by the higher cognitive demand associated with the GAME task performance than with the DR task performance.

A reward related increase from the rest level in θ power was observed in both tasks. The θ oscillation in the monkey medial prefrontal cortex is proposed to be related to the assessment of the reward [15]. The observed increase in θ power in this experiment may also be related to the assessment of reward or related to the attention to reward.

For both GAME and DR days during both the REST and TASK periods, the amount of information flows to aDMN from the two other networks and that to EN from pDMN was larger than the amount of the opposite direction of information flows (Figs. 3 and 4). Furthermore, the amount of information flow was significantly larger during the GAME task than during the REST period for the direction from EN to aDMN, that from pDMN to EN, and that from pDMN to aDMN, as compared with the opposite directions (Fig. 3). In all the three network areas, in relation to the GAME task performance, there was an increase in θ power, while in relation to the DR task performance, there was no significant change from the rest level (See Fig. 2). This may indicate that the GAME-task-related increase in θ power from the rest level in aDMN may be supported by more information flow conveyed by θ oscillation from EN and pDMN. The pDMN has a high baseline metabolic rate and is a tonically active region that continuously gathers information about the world [1]. It is known to be involved in integrating sensory information with memory [1]. In this study, we also observed high baseline θ activity in pDMN. Thus, pDMN may send the integrated information by θ oscillation to aDMN for cognitive control, directly or via EN.

The aDMN is reported to be important for social interaction [16]. It seems that aDMN plays the most important role during GAME by supporting social information processing associated with the GAME task performance.

In conclusion, our study indicates that DMN and EN are not anticorrelated, but work cooperatively, exchanging information by θ oscillation among the three networks for cognitive control of the game and DR task performance.

## Abbreviations

DMN: default mode network
aDMN: anterior DMN
pDMN: posterior DMN
EN: executive network
DR: delayed-response

## CRediT authorship Contribution statement

Tohru Kodama: Conceptualization, Formal analysis, Visualization, Writing - original draft,

Writing - review & editing

Takashi Kojima: Investigation, Formal analysis, Visualization

Yoshiko Honda: Resources, Visualization

Takayuki Hosokawa: Software

Akihiro Karashima: Software

Masataka Watanabe: Conceptualization, Methodology, Project administration, Visualization,

Writing - original draft, Writing - review & editing

## Data and code availability

Data will be made available on request.

## References

[1] M.E. Raichle, A.M. Leod, A.Z. Snyder, W.J. Powers, D.A. Gusnard, G.L. Shulman, A default mode of brain function, Proc. Natl. Acad. Sci. USA 98 (2) (2001) 676–682. doi: 10.1073/pnas.98.2.676.

[2] K. Koahino, Coactivation of default mode network and executive network regions in the human brain, in: M. Watanabe (Ed.), The Prefrontal Cortex as an Executive, Emotional, and Social Brain, Springer, Tokyo, 2017, pp. 247–276. DOI 10.1007/978-4-431-56508-6_13.

[3] J.F. Cavanagh, M.J. Frank, Frontal theta as a mechanism for cognitive control, Trends Cogn. Sci. 18 (8) (2014) 414–421. doi: 10.1016/j.tics.2014.04.012.

[4] G. Cona, F. Chiossi, S. Di Tomasso, G. Pellegrino, F. Piccione, P. Bisiacchi, G. Arcara, Theta and alpha oscillations as signatures of internal and external attention to delayed intentions: A magnetoencephalography (MEG) study, Neuroimage 205 (2020) 116295. doi: 10.1016/j.neuroimage.2019.116295.

[5] J.W.Y. Kam, J.J. Lin, A.K. Solbakk, T. Endestad, P.G. Larsson, R.T. Knight, Nat. Hum. Behav. 3(12) (2019) 1263–1270. doi: 10.1038/s41562-019-0717-0.

[6] T. Hosokawa, M. Watanabe, Prefrontal neurons represent winning and losing during competitive video shooting games between monkeys, J. Neurosci. 32 (22) (2012) 7662–7671. doi: 10.1523/JNEUROSCI.6479-11.2012.

[7] T. Kodama, T. Kojima, Y. Honda, T. Hosokawa, K.I. Tsutsui, M. Watanabe, Oral administration of methylphenidate (Ritalin) affects dopamine release differentially between the prefrontal cortex and striatum: A microdialysis study in the monkey, J. Neurosci. 37 (9) (2017) 2387–2394. doi: 10.1523/JNEUROSCI.2155-16.2017.

[8] A. Brovelli, M. Ding, A. Ledberg, Y. Chen, R. Nakamura, and S. L. Bressler, Beta oscillations in a large-scale sensorimotor cortical network: Directional influences revealed by Granger causality, Proc. Natl. Acad. Sci. USA 101 (26) (2004) 9849–9854. DOI: 10.1073/pnas.0308538101

[9] A.K. Seth, A MATLAB toolbox for Granger causal connectivity analysis, J. Neurosci. Methods 186 (2) (2010) 262–273. doi: 10.1016/j.jneumeth.2009.11.020.

[10] T. H. Ryan, Significance tests for multiple comparisons of proportions, variances, and other statistics, Psychol. Bull. 57 (1960) 318–328. https://doi.org/10.1037/h0044320.

[11] T. Kojima, H. Onoe, K. Hikosaka, K. Tsutsu, H. Tsukada, M. Watanabe, Default mode of brain activity demonstrated by positron emission tomography imaging in awake monkeys: higher Rest-related than working memory-related activity in medial cortical areas, J. Neurosci. 29 (46) (2009) 14463–14471. doi: 10.1523/JNEUROSCI.1786-09.2009.

[12] R. Scheeringa, M.C. Bastiaansen, K.M. Petersson, R. Oostenveld, D.G. Norris, P. Hagoort, Frontal θ EEG activity correlates negatively with the default mode network in Resting state, Int. J. Psychophysiol. 67 (3) (2008) 242–251. doi: 10.1016/j.ijpsycho.2007.05.017.

[13] P. Sauseng, J. Hoppe, W. Klimesch, C. Gerloff, F.C. Hummel, Dissociation of sustained attention from central executive functions: Local activity and interregional connectivity in the theta range, Euro. J. Neurosci. 25 (2), (2007) 587–593. doi: 10.1111/j.1460-9568.2006.05286.x.

[14] R. Scheeringa, K.M. Petersson, R. Oostenveld, D.G. Norris, P. Hagoort, M.C.M. Bastiaansen, Trial-by-trial coupling between EEG and BOLD identifies networks related to alpha and theta EEG power increases during working memory maintenance, Neuroimage 44 (3) 1224–1238. doi: 10.1016/j.neuroimage.2008.08.041.

[15] T. Tsujimoto, H. Shimazu, Y. Isomura, K. Sasaki, Theta oscillations in primate prefrontal and anterior cingulate cortices in forewarned reaction time Tasks, J. Neurophysiol. 103 (2) (2010) 827–843. doi: 10.1152/jn.00358.2009.

[16] D.M. Amodio, C.D. Frith, Meeting of minds: the medial frontal cortex and social cognition, Nat. Rev. Neurosci. 7(4) (2006) 268–277. doi: 10.1038/nrn1884.

